# Pushing the limits of *de novo* genome assembly for complex prokaryotic genomes harboring very long, near identical repeats

**DOI:** 10.1101/300186

**Authors:** Michael Schmid, Daniel Frei, Andrea Patrignani, Ralph Schlapbach, Jürg E. Frey, Mitja N.P. Remus-Emsermann, Christian H. Ahrens

**Affiliations:** Agroscope, Research Group Molecular Diagnostics, Genomics & Bioinformatics, CH-8820 Wädenswil, Switzerland; SIB Swiss Institute of Bioinformatics, CH-8820 Wädenswil, Switzerland; Functional Genomics Center Zurich, CH-8057 Zurich, Switzerland; School of Biological Sciences, University of Canterbury, 8140 Christchurch, New Zealand; Biomolecular Interaction Centre, University of Canterbury, 8140 Christchurch, New Zealand

**Keywords:** *De novo* genome assembly, repeat-rich genomes, complete genome, prokaryotes, whole genome sequencing, *Pseudomonas*, shufflon, Pacific Biosciences, Oxford Nanopore Technologies

## Abstract

Generating a complete, *de novo* genome assembly for prokaryotes is often considered a solved problem. However, we here show that *Pseudomonas koreensis* P19E3 harbors multiple, near identical repeat pairs up to 70 kilobase pairs in length. Beyond long repeats, the P19E3 assembly was further complicated by a shufflon region. Its complex genome could not be *de novo* assembled with long reads produced by Pacific Biosciences’ technology, but required very long reads from the Oxford Nanopore Technology. Another important factor for a full genomic resolution was the choice of assembly algorithm.

Importantly, a repeat analysis indicated that very complex bacterial genomes represent a general phenomenon beyond *Pseudomonas*. Roughly 10% of 9331 complete bacterial and a handful of 293 complete archaeal genomes represented this dark matter for *de novo* genome assembly of prokaryotes. Several of these dark matter genome assemblies contained repeats far beyond the resolution of the sequencing technology employed and likely contain errors, other genomes were closed employing labor-intense steps like cosmid libraries, primer walking or optical mapping. Using very long sequencing reads in combination with assemblers capable of resolving long, near identical repeats will bring most prokaryotic genomes within reach of fast and complete *de novo* genome assembly.

## Introduction

The enormous pace in next generation sequencing (NGS) technology development (1) has led to an exponential increase in the number of publicly available, complete prokaryotic genome assemblies (2). Due to their lower complexity, the genome assembly problem, mainly caused by the presence of long repeats, had been considered solved for prokaryotes. However, despite advances from Pacific Biosciences (PacBio) and more recently Oxford Nanopore Technologies (ONT) to sequence very long reads (above 10kb and well beyond) which allow *de novo* bacterial genome assembly (3), the percentage of complete genomes is still low compared to the large number of fragmented assemblies based on Illumina short reads (2, 4), most of which remain at a permanent draft stage. This can in part be attributed to a considerable lag between the development of a new technology and its broader adoption, and to the higher costs of PacBio and ONT. The fragmented Illumina-based genome assemblies can represent serious limitations for follow up analyses (5–7). We and many others could recently illustrate the benefits of relying on complete genomes, e.g., for comparative genomics, where a study of repeat-rich genomes indicated that even core genes could be missed on top of accessory genes, and for metagenomic studies, which benefited from the availability of complete genomes as a larger fraction of reads could be assigned down to the strain level (8). Moreover, complete genome sequences represent the optimal basis for accurate genome annotation (9), to create accurate, genome-scale metabolic models (10), to track the spread of mobile genetic elements including those carrying antimicrobial resistance genes (4), to explore patterns for the emergence of drug resistance (11) and as the optimal starting point for functional genomics studies (5). The development of sophisticated assembly algorithms (12–14) has and will continue to increase the percentage of complete genomes. Several large genome sequencing initiatives for prokaryotes will benefit from these advances. These include the Genomic Encyclopedia of Bacteria and Archaea (GEBA) initiative, which aims to sequence the type strain of ∼11,000 species with a validly published name (15) and the NCTC 3000 project, which aims to completely sequence reference bacteria and viruses of public health importance (http://www.sanger.ac.uk/resources/downloads/bacteria/nctc/).

Furthermore, efforts to isolate individual strains from the complex soil or rhizosphere microbiomes which synthesize urgently needed, novel antibiotics (16) or carry out promising biological functions, e.g., the protection of crops from pathogens (17, 18), are gaining momentum. The genus *Pseudomonas*, which belongs to the gammaproteobacteria, is of particular interest as it includes several pathogens like *Pseudomonas aeruginosa* and *Pseudomonas syringae*, but also species with a potential use as biocontrol agents (17, 18) or for bio-remediation (19). *P. koreensis* was first isolated from Korean agricultural soil (20). It too exhibited antagonistic activities: a biosurfactant was active both against *Phytium ultimum* in a tomato model system (21) and against *Phytophthora infestans* infecting potato plants (22). Another *P. koreensis* strain, CRS05-R05, was isolated from the rice rhizosphere and showed biocontrol activity against the rice weevil *Sitophilus oryzae* (23), a pest of stored foods.

Challenged by the complexity of the *Pseudomonas koreensis* isolate (P19E3) genome, we here push the limits of *de novo* genome assembly for a very complex bacterial genome. We exclusively relied on sequencing data without labor-intense steps like cosmid or BAC libraries, optical mapping or primer-walking that otherwise would be required to scaffold and close gaps of highly complex prokaryotic genomes. The long, near identical repeats of strain P19E3 could only be resolved by very long ONT reads. Importantly, the analysis of the repeat complexity of 9331 publicly available complete bacterial and 293 archaeal genomes, which we release here, established that roughly 10% of genomes represented this “dark matter” of prokaryotic *de novo* genome assembly (24), i.e., genomes that are particularly difficult to assemble, either due to the presence of several hundred repeats (ca. 7%), or very long highly similar repeats (ca. 3%).

## Methods

### Repeat analysis for genus *Pseudomonas* and all completely sequenced prokaryotes

All publicly available, completely sequenced genomes of the genus *Pseudomonas* (a total of 270) were downloaded from the National Center of Biotechnology Information’s (NCBI) GenBank (Feb. 8, 2018), i.e. NCBI assembly level “Complete genome”, the highest level compared to levels “Chromosome”, “Contigs”, and “Scaffold”. A repeat analysis was carried out as described earlier (8), allowing us to triage the genomes into 3 genome assembly complexity classes (25) and to visualize the data. The same analysis was performed for all complete prokaryotic genomes from GenBank (9331 bacteria and 293 archaea, Feb. 23, 2018).

### Bacterial strain, genomic DNA extraction & sequencing

*Pseudomonas koreensis* P19E3 was isolated from healthy marjoram (*Origanum marjorana*) leaf material during an isolation survey on an organic herb farm (Boppelsen, Switzerland) in summer 2014 and classified by matrix-assisted laser desorption/ionization time of flight (MALDI-TOF) analysis (26). Genomic DNA (gDNA) was isolated (Sigma GenElute kit), and PacBio SMRT sequencing was performed on an RS II machine (4 SMRT cells, P6-C4 chemistry) aiming to *de novo* assemble a complete genome as described (9). A size selection was performed with the BluePippin system, first for fragments larger than 10 kb (2 SMRT cells), later for fragments larger 20 kb (2 SMRT cells). For ONT sequencing, high molecular weight gDNA was extracted according to a recent protocol (27). The ONT libraries were prepared using a 1D^2^ sequencing kit (SQK-LSK308) and sequenced on two R9.5 flow cells (FLO-MIN107). Base calling was performed using Albacore v.2.1.3 (community.nanoporetech.com/downloads). Finally, a 2 × 300 base pair (bp) paired end library was prepared using Illumina’s Nextera XT DNA kit and sequenced on a MiSeq. ONT, PacBio and Illumina data were uploaded to NCBI SRA and can be accessed via BioProject PRJNA436895.

### Genome assembly

#### Assemblies based on PacBio data

PacBio reads were assembled, once with HGAP v.3 (12) and once with Flye v.2.3 (14). HGAP3 was run on SMRT Analysis v.2.3.0 and Protocol “RS_HGAP_Assembly.3” with data of all 4 SMRT cells (default parameters, except: min. subread length: 500; min. polymerase read quality: 0.80; estimated genome size: 7.5 Mb). To assemble the data with Flye, we first extracted the subreads using Protocol “RS_Subreads.1” of SMRT Analysis (min. subread length: 500, min. polymerase read quality: 0.83). The resulting subread FASTA files were then processed using Flye v.2.3 applying standard parameters and an estimated genome size of 7.5 Mb. No further polishing was performed for these two assemblies on top of the polishing and correction steps included in the assembly pipelines.

#### Flye assembly of Oxford Nanopore data

Base called ONT reads were filtered (30 kb or longer), and assembled with Flye v.2.3 (14) using standard parameters and performing three polishing iterations on Flye’s polishing stage (-i 3 --nano-raw [ont_reads.fastq] -g 7.7m). The genome size was estimated by a first, explorative Flye assembly, before removing a few spurious contigs, i.e., short contigs mainly consisting of repeated short DNA motifs and with a low coverage. The resulting circular contigs were linearized (start of *dnaA* gene for chromosome, putative intergenic region for plasmids). Also, a repeat analysis was performed to ensure that repeats longer than 10 kb were not disrupted when linearizing the circular chromosome and plasmids.

#### Assembly of putative small plasmids

To catch potential small plasmids which would be missed in the Flye assembly due to size selection for long reads, we assembled the MiSeq 2 × 300 bp data using plasmidSPAdes (SPAdes v.3.11.1) (28).

#### Error correction & polishing of the ONT Flye assembly

The post-processed ONT draft assembly was subsequently corrected by three iterative Racon runs (29). For each run, the filtered ONT reads were first mapped to the draft assembly using GraphMap v.0.5.2 (30). Reads entirely contained in long genomic repeats (longer than 10 kb) were excluded from mapping using samtools v.1.6 (31). The resulting SAM file was then provided to Racon v.0.5.0 for correction of the assembly. A polishing step using the ONT data was performed applying Nanopolish v.0.8.5 (methylation aware mode “-methylation-aware=dcm,dam”) (32). Like before, reads which were entirely contained in a repeat region were discarded for polishing. To correct any remaining small assembly errors, a last polishing step was performed using the 2 × 300 bp Illumina data by mapping the raw reads to the assembly (bwa mem v.0.7.12) and using FreeBayes v.1.1.0 (https://arxiv.org/abs/1207.3907v2). As final step, a repetitive rearrangement region was manually corrected (see below).

#### Comparison of different assembly stages and check of final assembly

The different assembly stages were compared to the final Illumina-polished assembly using Quast v.4.6.3 (33). A last validation of the final assembly with respect to potential large scale mis-assemblies was performed using Sniffles v.1.0.8 (34) based on a mapping of ONT data with the NGM-LR mapper v.0.2.6 (34). The ONT mapping was subsequently visually inspected using the Integrated Genome Viewer (IGV) (35). Potential small-scale errors were checked using Illumina data and FreeBayes in combination with visual inspection of the mapping. An overview of the steps carried out for genome assembly, polishing and comparison of different assemblies is shown as workflow in Supplementary Figure S1.

#### Local hybrid assembly of a highly repetitive rearrangement region

A highly repetitive – and by the Flye assembler mis-assembled – rearrangement region on plasmid 2 of the polished assembly was resolved using a hybrid assembly strategy applying ONT and Illumina data. First, all ONT reads spanning the region and covering about 10 kb of unique regions on both sides were extracted from a previous mapping with GraphMap (see above). Second, proovread v.2.14.1 (36) was used to polish the ONT reads with Illumina data. Two specific ONT reads which could be polished by proovread (without apparent dips in Illumina data coverage) were selected and again processed with proovread and all Illumina reads. Both polished ONT reads were cut at the same unique positions up- and downstream of the rearrangement region and compared using progressiveMauve v.2.3.1 (37). One of the polished reads was then used to replace the mis-assembled region of the polished assembly of plasmid 2. Next, the inverted repeats on the rearrangement region were identified and a multiple sequence alignment was created using MUSCLE v.3.8 (38) and visualized with WebLogo (39).

#### Repeat analysis for ONT assembly and comparison to the PacBio assemblies

A repeat analysis was performed for the final ONT assembly as described earlier (25), only considering repeats longer than 30 kb. The output was simplified and formatted as BED file to enable visualization as a circular plot. Nucmer 3.1 (40) was used to compare and map the PacBio assemblies (as query) onto the final ONT assembly (serving as reference) (parameters: “--mum --simplify -b 5000 -D 30 -d 0.25”). Matches smaller than 20 kb and below 95% identity were removed. The output was again formatted as BED file. The final ONT assembly, results from its repeat analysis and the mapping of PacBio assemblies were combined and plotted using Circos v.0.69 (41).

#### Phylogenetic analysis

GenBank records of selected, completely sequenced *Pseudomonas* strains were downloaded from NCBI RefSeq (March, 2018). The predicted CDS of 107 known housekeeping genes (42) were extracted from 18 strains (including *P. koreensis* P19E3, and *Azotobacter vinelandii DJ* as outgroup) and used to calculate a maximum likelihood phylogenetic tree using bcgTree (43), as described previously (8).

#### Genome annotation

The genome sequence of *P. koreensis* P19E3 was submitted to NCBI GenBank and annotated with the genome annotation pipeline for prokaryotes (PGAP) (44). It is available under GenBank accession numbers CP027477 to CP027481.

## Results

### Repeat analysis for the genus *Pseudomonas*

A bacterial strain isolated during a screening of herbal plants for food spoiling and pathogenic bacteria was assigned to the species *Pseudomonas koreensis* by MALDI biotyping (26). However, although PacBio long read sequencing was used, it was not possible to assemble the genome *de novo* into a single complete chromosome.

Therefore, we explored the predicted genome assembly complexity (25) for the genus *Pseudomonas*. Koren and colleagues had classified bacterial genomes in three categories based on i) the number of repeats greater than 500 bp with a nucleic acid sequence similarity higher than 95%, and ii) the length of the longest repeat above 95% similarity: class I genomes contain less than 100 repeats of up to 7 kb (approximate length of the rDNA operon), class II genomes with more than 100 repeats of up to 7kb, and finally class III genomes with repeats that can be substantially longer than the 7 kb of the rDNA operon. We analyzed 270 publicly available complete *Pseudomonas* genome sequences (see Methods), which included two *P. koreensis* strains (Figure 1). When selecting a plot range identical to that used by Koren and colleagues, 245 genomes with repeat lengths of up to 30 kb and up to 300 repeats of 500 bp or above were captured (Figure 1A).

**Figure 1.**
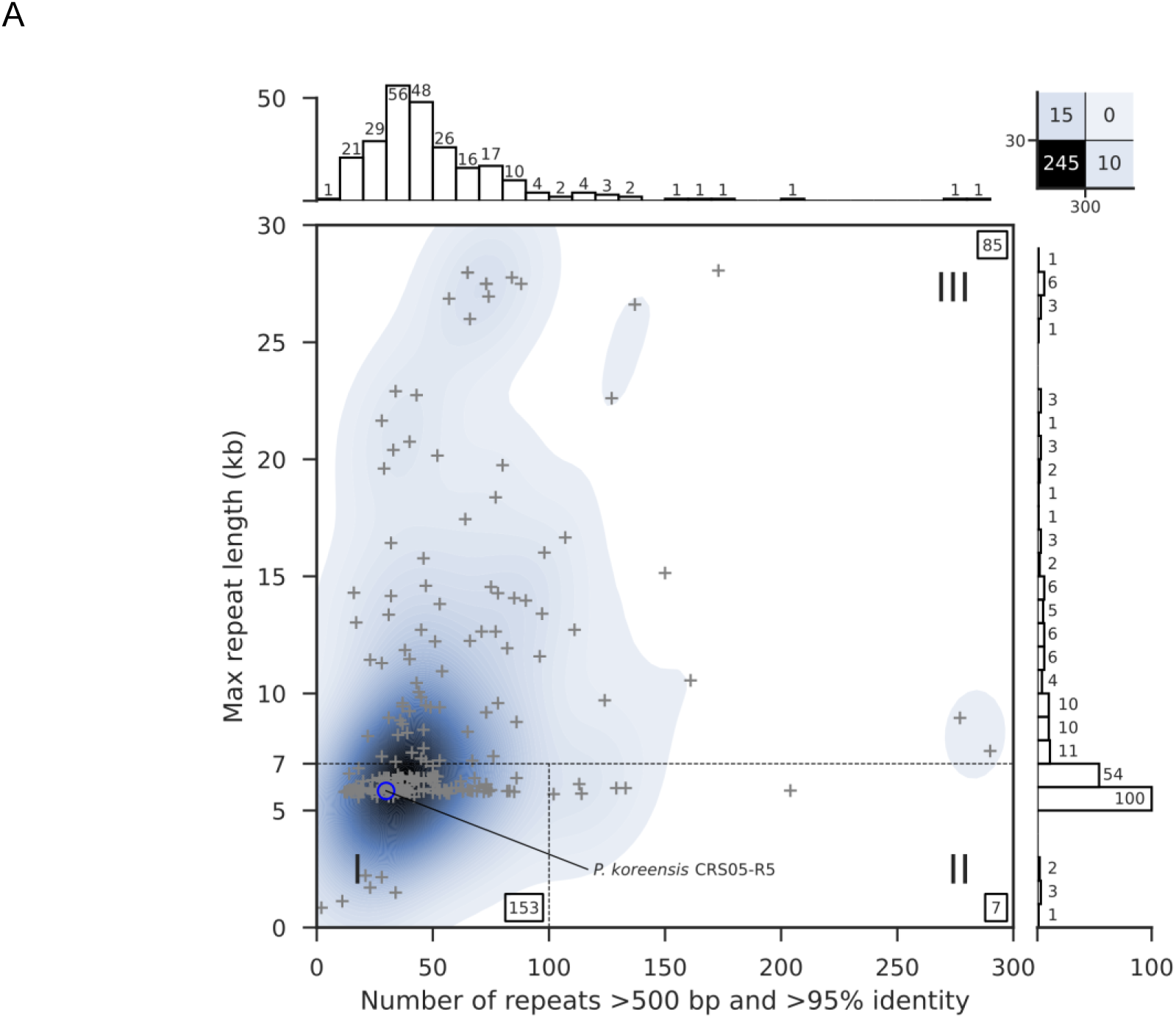
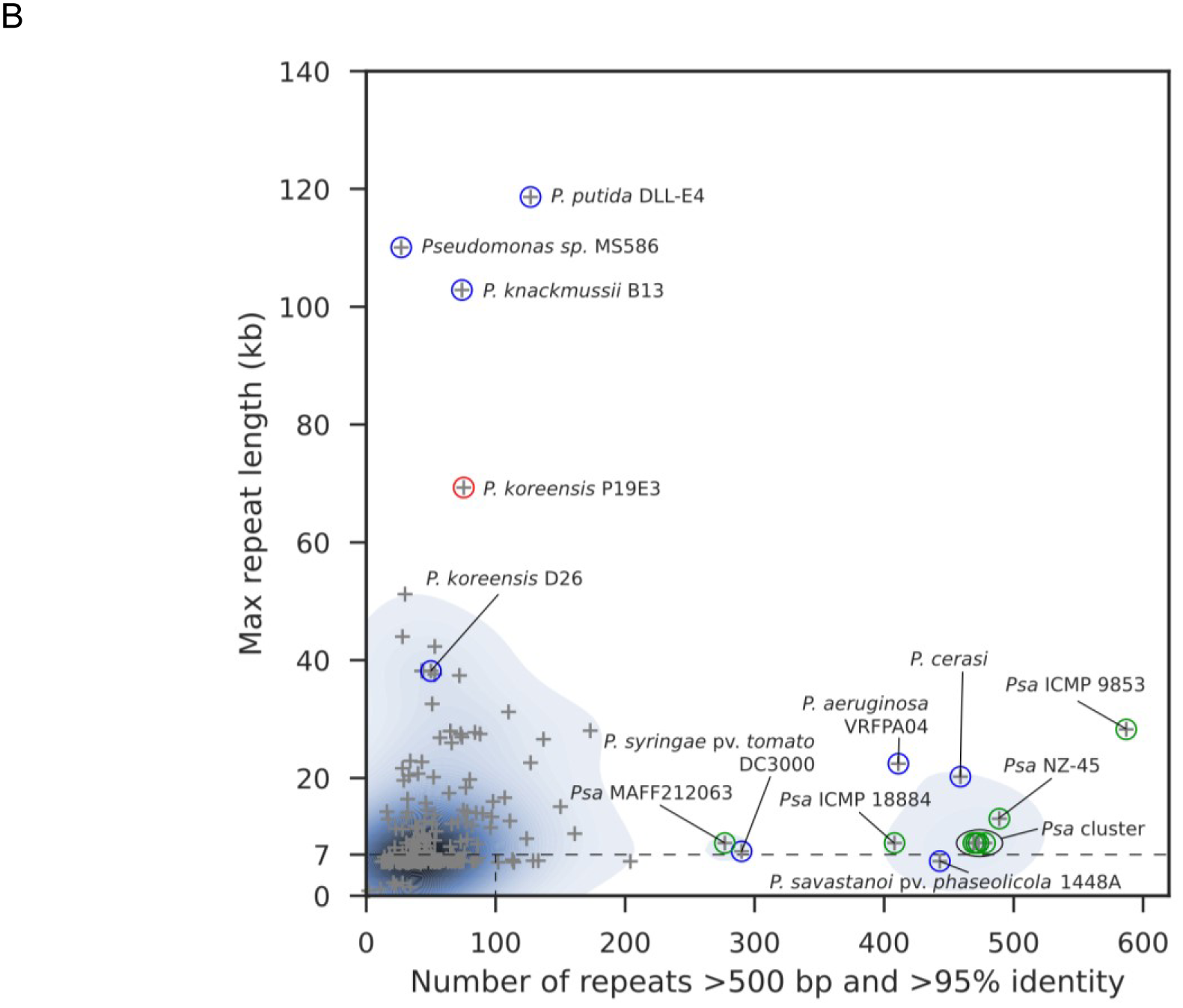
Genome assembly complexity of the genus *Pseudomonas*. **(A)** Original representation of genome assembly complexity, as described by Koren and colleagues (25). The highlighted *P. koreensis* strain CRS05-R5 strain represents a class I genome. A total of 25 genomes are not covered in this plot, including 10 genomes with more than 300 repeats and 15 genomes whose longest repeats exceeded 30 kb (upper right box). **(B)** Overview of the most complex Pseudomonas strains. *P. koreensis* strain P19E3 is labeled (red circle) and several other strains with complex genomes, including 8 *Psa* strains (green).

Our repeat analysis indicated that 15 *Pseudomonas* genomes had longer repeats and 10 exhibited a higher overall number of repeats (see upper right box, Figure 1A), corresponding to roughly 9% of the *Pseudomonas* strains analyzed. When extending the plot range (Figure 1B), we captured these highly complex genomes as well. Of note was a cluster of 10 *Pseudomonas* genomes with more than 400 repeats (Figure 1B, Supplementary Table S1), 8 of which represented different *P. syringae* pathovar (*pv.*) *actinidiae* (*Psa*) strains, the causal agent of Kiwi fruit plant canker (45). The only completely sequenced *Psa* strain not covered in this extended range (more than 300 repeats) was strain MAFF212063 that harbored 277 repeats (Figure 1B). Among the most complex genomes in terms of repeat length were many *P. aeruginosa* strains (11 of 15), *P. koreensis* strain 26 (with repeats of 38 kb), *P. knackmussii* strain B13 with repeats of around 103 kb and *P. putida* DLL-E4 with repeats close to 120 kb (Figure 1B, Table 1). These strains also included our *P. koreensis* strain P19E3, which harbored near identical repeats of almost 70 kb (red circle, Figure 1B). The steps that finally enabled us to *de novo* assemble its complete genome purely by using long sequencing reads are described in more detail in the sections below.

**Table 1.** Overview of the most complex *Pseudomonas* genomes. 15 genomes are shown whose longest repeat pair was longer than 30 kb (sequence identity above 95%). Pseudomonas koreensis P19E9, *de novo* assembled in this study is shown in bold.

| Species | Strain | Accession(s) | Longest repeat pair (bp) | Number of repeats |
| --- | --- | --- | --- | --- |
| *Pseudomonas putida* | DLL-E4 | CP007620 | 118,617 | 127 |
| *Pseudomonas sp.* | MS586 | CP014205 | 110,079 | 27 |
| *Pseudomonas knackmussii* | B13 | HG322950 | 102,855 | 74 |
| ***Pseudomonas koreensis*** | **P13E9** | **CP027477 to CP027481** | **68,866** | **74** |
| *Pseudomonas aeruginosa* | PB353 | CP025051, CP025052 | 51,221 | 30 |
| *Pseudomonas aeruginosa* | PA1R | CP004055 | 43,998 | 28 |
| *Pseudomonas aeruginosa* | PA11803 | CP015003 | 42,317 | 53 |
| *Pseudomonas aeruginosa* | PA_D5 | CP012579 | 38,173 | 43 |
| *Pseudomonas aeruginosa* | PA_D25 | CP012584 | 38,172 | 44 |
| *Pseudomonas aeruginosa* | PA_D22 | CP012583 | 38,171 | 43 |
| *Pseudomonas aeruginosa* | PA_D16 | CP012581 | 38,171 | 43 |
| *Pseudomonas koreensis* | D26 | CP014947 | 38,167 | 50 |
| *Pseudomonas aeruginosa* | DN1 | CP017099, CP018048 | 37,573 | 53 |
| *Pseudomonas aeruginosa* | NCGM257 | AP014651 | 37,397 | 72 |
| *Pseudomonas aeruginosa* | PA7790 | CP014999, CP015000 | 32,594 | 51 |
| *Pseudomonas aeruginosa* | Carb01 63 | CP011317 | 31,243 | 110 |

### Genome assembly of *P. koreensis* P19E3

Due to the genome complexity classification of previously sequenced *P. koreensis* strains, where one strain had a complex class III genome with repeats of around 38 kb (Table 1), we first attempted to sequence and *de novo* assemble the genome using PacBio’s long read technology (46) combined with a size selection step (inserts greater 10 kb) and two SMRT cells, a strategy that had proven useful for a class III genome before (9). However, an assembly with HGAP3 resulted in 25 contigs and could not resolve the chromosome and plasmids; presumably due to the presence of long repeat sequences.

Adding data from two more SMRT cells, this time relying on an even longer BluePippin size selection cut-off (greater 20 kb), improved the assembly; it returned 14, i.e. fewer contigs, but did not allow us to generate a complete genome assembly.

To assess the possibility that HGAP3 had difficulties in resolving long, near identical repeats, we then explored the Flye assembler (14), which was specifically designed to address this issue. The assembly created by Flye reduced the number of contigs to 7 and also showed less redundancy in repeat regions compared to the HGAP3 assembly (Figure 3). However, it did not result in a completely gapless assembly either.

**Figure 3.**
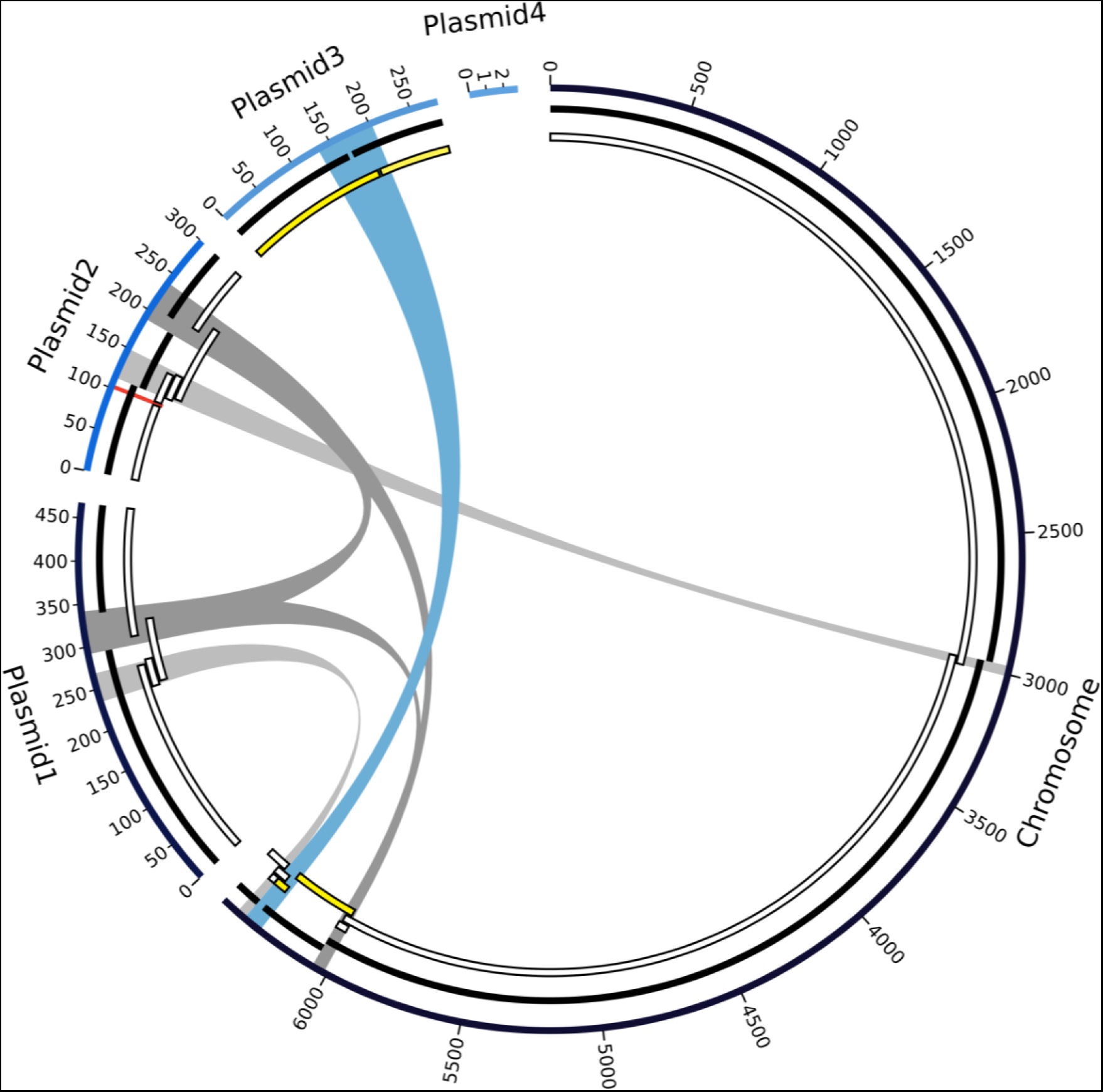
Graph of the final high-quality *P. koreensis* P19E3 genome assembly. For display, the five circular elements (one chromosome, 4 plasmids) were linearized and not drawn to scale. Going from outward to inward circles: 1) ONT data assembly using Flye, 2) PacBio data assembled with Flye and 3) PacBio data assembled with HGAP3 are shown. Repeats above 30 kb are shown in the center (blue and gray bands, blue showing the longest repeat), which are also listed in Table 4; they coincide with areas where the PacBio-based assemblies were fragmented. A genomic region identified as a structural variation (ca. 5 kb) is also shown (red mark on plasmid 2). For the HGAP3 assembly with PacBio reads (innermost circle), regions are marked (in yellow) which, due to an assembly error, mapped to both chromosome and plasmid; Flye was able to resolve this region. Since plasmid 4 did not get assembled with the long read technologies, due to its short size, there is no PacBio counterpart in the PacBio tracks.

To create a gapless complete genome assembly, we therefore ended up sequencing high molecular weight gDNA (see Methods) on the ONT MinION platform, relying on reads from two ONT 1D^2^ libraries (Table 2). With an N50 read length of above 44 kb (after having filtered for reads longer 30 kb), the read length of these two runs was significantly longer than the average N50 of close to 17 kb from the previous PacBio runs (Table 2, Figure 2).

**Figure 2.**
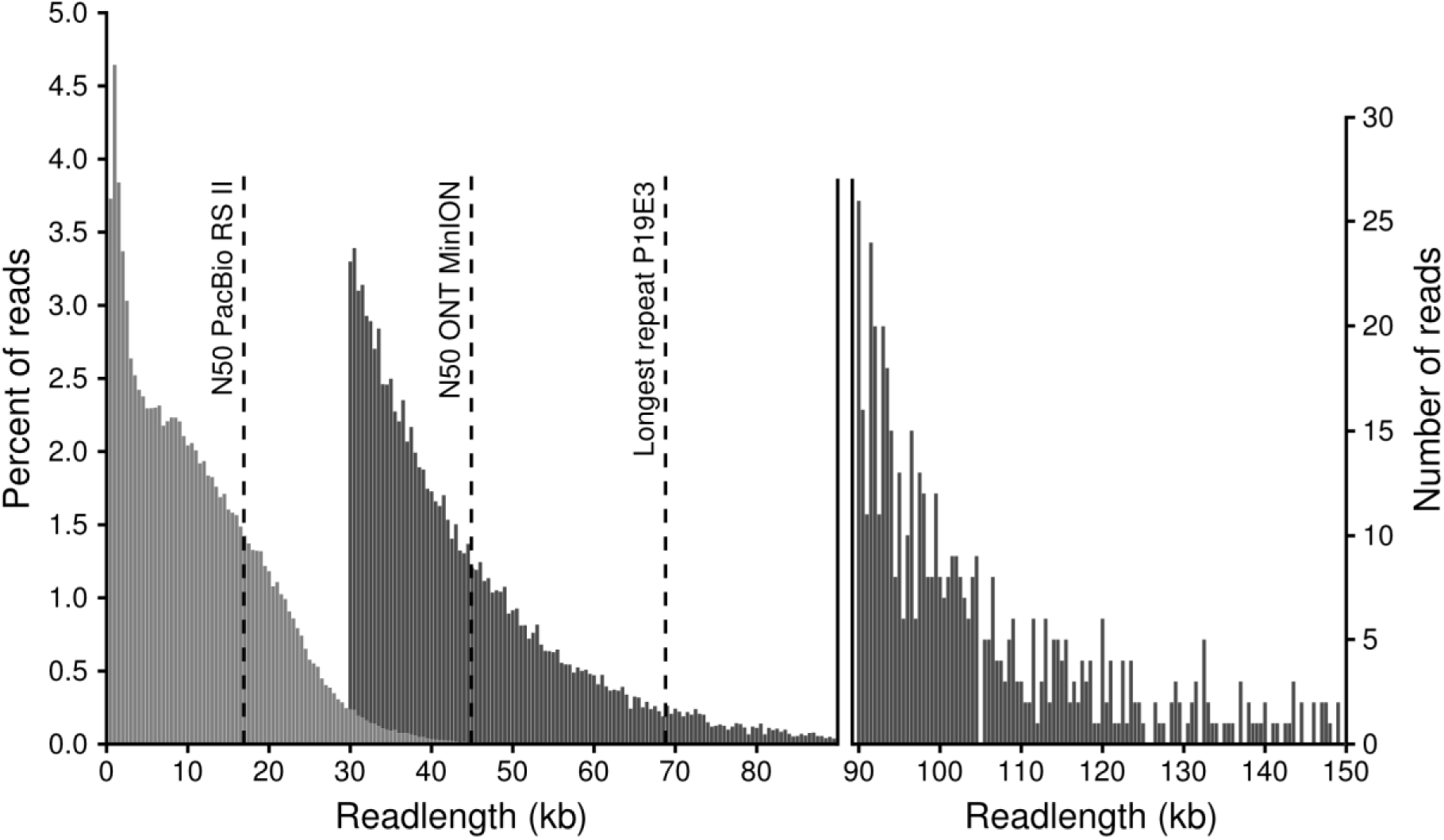
Length distribution of PacBio (RSII platform with BluePippin size selection, light gray) and ONT reads (MinION 1D^2^ libraries, dark gray). ONT can generate very long reads; we here only show reads above 30 kb, as shorter reads were not used for the assembly. The percentage of reads per length bin (500 bp) is shown on the left, while the number of ONT reads longer than 90 kb is shown in the right legend. Reads longer than 150 kb (115 reads in total, longest completely mappable read 288 kb) are not shown. The three dashed lines show (from left to right) the N50 read length of PacBio subreads, the N50 read length of ONT reads above 30 kb and the length of the longest repeat in the *P. koreensis* P19E3 genome.

**Table 2.** Basic statistics of data from three applied sequencing technologies.

| Technology | Number of units sequenced | Number of reads | N50 read length | Total amount of data in base pairs |
| --- | --- | --- | --- | --- |
| Illumina, MiSeq | Part of one flow cell (300 bp paired-end) | 4,471,800 (read pairs) | 242 bp | 753 Mb |
| PacBio, RS II <sup>*1</sup> | Four SMRT cells (P6-C4 chemistry) | 239,198 | 16.9 kb | 2.7 Gb |
| ONT, MinION <sup>*2</sup> | Two flow cells (R9.5 cells, 1D<sup>2</sup> library) | 34,167 | 44.9 kb | 1.6 Gb |

<sup>\*1</sup> Statistics based on extracted subreads. After length (> 500 bp) and quality (0.83) filtering.
<sup>\*2</sup> Statistics after filtering for reads longer than 30 kb.

A final *de novo* genome assembly attempt based on ONT MinION data resulted in one chromosome (6.44 Mb) and three large plasmids (468 kb, 300 kb, 283 kb) with a coverage exceeding 150-fold (Supplementary Table S3). In addition, a small plasmid of 2.8 kb size was identified by using plasmidSPAdes (28) (Figure 3, Table 3). Illumina data (∼100-fold coverage) was mainly used for correction of potential single base-pair errors (47) and to achieve a very low final error rate (3, 4). The various steps carried out for assembly, polishing and comparison of different assemblies are shown in Figure S1. A multilocus sequence analysis based on the comparison of 107 conserved housekeeping genes (42) allowed to establish the phylogenetic relationship of our strain with selected *Pseudomonas* strains and confirmed the MALDI-TOF assignment as a *P. koreensis* strain (Supplementary Figure S2).

**Table 3.**
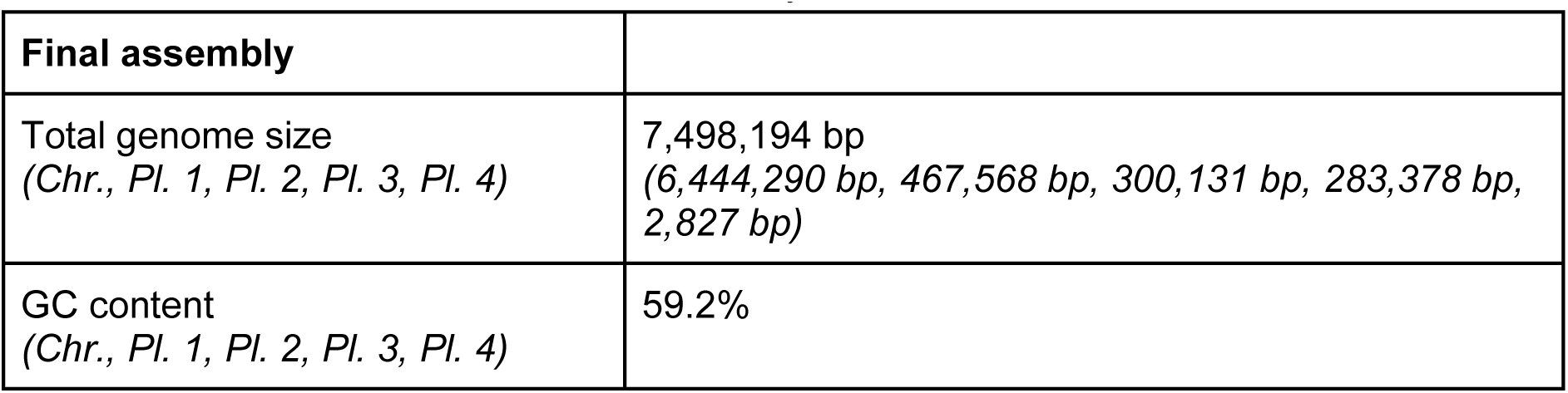
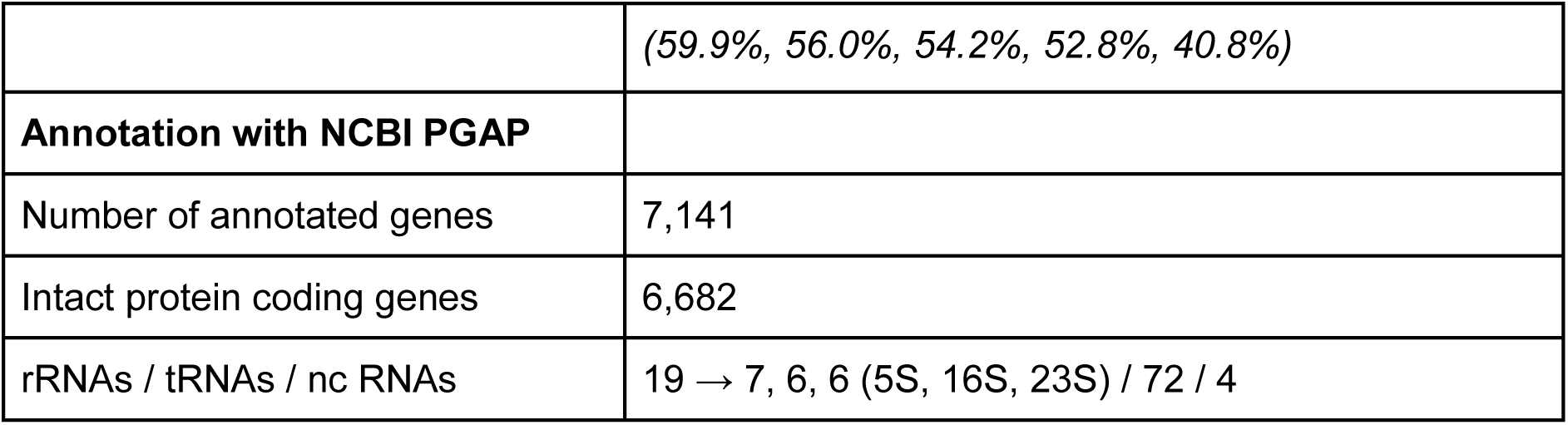
Basic metrics about the final assembly and its annotation.

| Final assembly |  |
| --- | --- |
| Total genome size<br>(Chr., Pl. 1, Pl. 2, Pl. 3, Pl. 4) | 7,498,194 bp<br>(6,444,290 bp, 467,568 bp, 300,131 bp, 283,378 bp, 2,827 bp) |
| GC content<br>(Chr., Pl. 1, Pl. 2, Pl. 3, Pl. 4) | 59.2% |
|  | (59.9%, 56.0%, 54.2%, 52.8%, 40.8%) |
| **Annotation with NCBI PGAP** |  |
| Number of annotated genes | 7,141 |
| Intact protein coding genes | 6,682 |
| rRNAs / tRNAs / nc RNAs | 19 → 7, 6, 6 (5S, 16S, 23S) / 72 / 4 |

We also analyzed the repeat complexity of the final genome assembly and identified three repeat pairs and one repeat triplet when filtering for repeats longer than 30 kb (Table 4). All repeats had a sequence identity higher than 99.7% with few gaps in the respective blastn alignment of repeat pairs. The longest repeat pair (69 kb) was shared between the chromosome and plasmid 3 (Figure 3). This repeat pair could not be resolved when relying solely on PacBio data. The second longest repeat was a triplet of 46 kb located on the chromosome, on plasmid 1, and on plasmid 2, which could not be resolved either using only PacBio data. The same held true for the third longest repeat (37 kb) located on the chromosome and on plasmid 2. In contrast to HGAP3, a fourth repeat pair of 34 kb length could be resolved by the Flye assembler using only PacBio data. Of note, one copy of this fourth repeat pair was directly adjacent to the longest repeat on the copy located on the chromosome, potentially further complicating this assembly. A putative shufflon recombination site on plasmid 2 (see below, and Figure 3, red mark), which had not been spanned by the HGAP3 PacBio assembly, was also resolved by the Flye PacBio assembly.

**Table 4.** Length and similarity of all repeats pairs/groups longer than 30 kb

| Name | Length | Identity | Alignment gaps in bp | Genomic elements involved |
| --- | --- | --- | --- | --- |
| Repeat 1 | 69 kb | 99.7 % | 131 bp | Chromosome, Plasmid 3 |
| Repeat 2* | 46 kb | 99.8 % | 100 bp | Chromosome, Plasmid 1, Plasmid 2 |
| Repeat 3 | 37 kb | 99.8 % | 58 bp | Chromosome, Plasmid 2 |
| Repeat 4 | 34 kb | 99.9 % | 35 bp | Chromosome, Plasmid 1 |

\* For repeat 2, the average identity and number of gaps of all three possible comparisons is given.

### Polishing and quality assessment of the final assembly

Error correction and polishing of the P19E3 genome was done in several stages (Supplementary Figure S1; for software and respective versions used, see Methods). As the Flye assembler does not use quality value information, we first carried out three rounds of error correction with the ONT data using Racon (29), which can employ quality information encoded in FASTQ files. Compared to the initial Flye assembly, this step reduced the number of indel and mismatch assembly errors (Supplementary Table S2). Another step using Nanopolish (32) further reduced their amount. Remaining errors (compared to the final assembly), i.e. mostly single base-pair errors, were removed in a final polishing step using Illumina MiSeq data.

We next mapped the ONT and Illumina data back against the polished genome assembly to assess if small scale mis-assemblies could be detected, combining an automated detection using Freebayes and a visual inspection with IGV. No smaller scale mis-assemblies were detected. To find evidence for potential larger scale errors like inversions, we first mapped the ONT data on the final assembly using NGM-LR and then ran Sniffles on the mapping. However, except for a complex rearrangement region which was resolved later (see below), no evidence for a mis-assembly was found. The mapping statistics for all sequencing technologies are listed in Supplementary Table S3.

### Resolving a highly repetitive rearrangement region

During the error correction of the ONT assembly, we identified a region of about 5 kb on plasmid 2, which stood out due to the poor mappability of both ONT and Illumina reads (Positions 99,590 to 104,500). A tblastx search indicated that this region contained a shufflon system (48) (Figure 4A). To resolve this putative shufflon region, a tailored hybrid polishing strategy using a combination of ONT and Illumina data from this region (see Methods) and the software proovread (36) was applied, which allowed to uncover evidence for two major variants of the shufflon (Figure 4 A,B). Since a mapping of ONT and Illumina data did not indicate an abundance difference of the two variants, one variant was picked to represent the shufflon in the final assembly.

**Figure 4:**
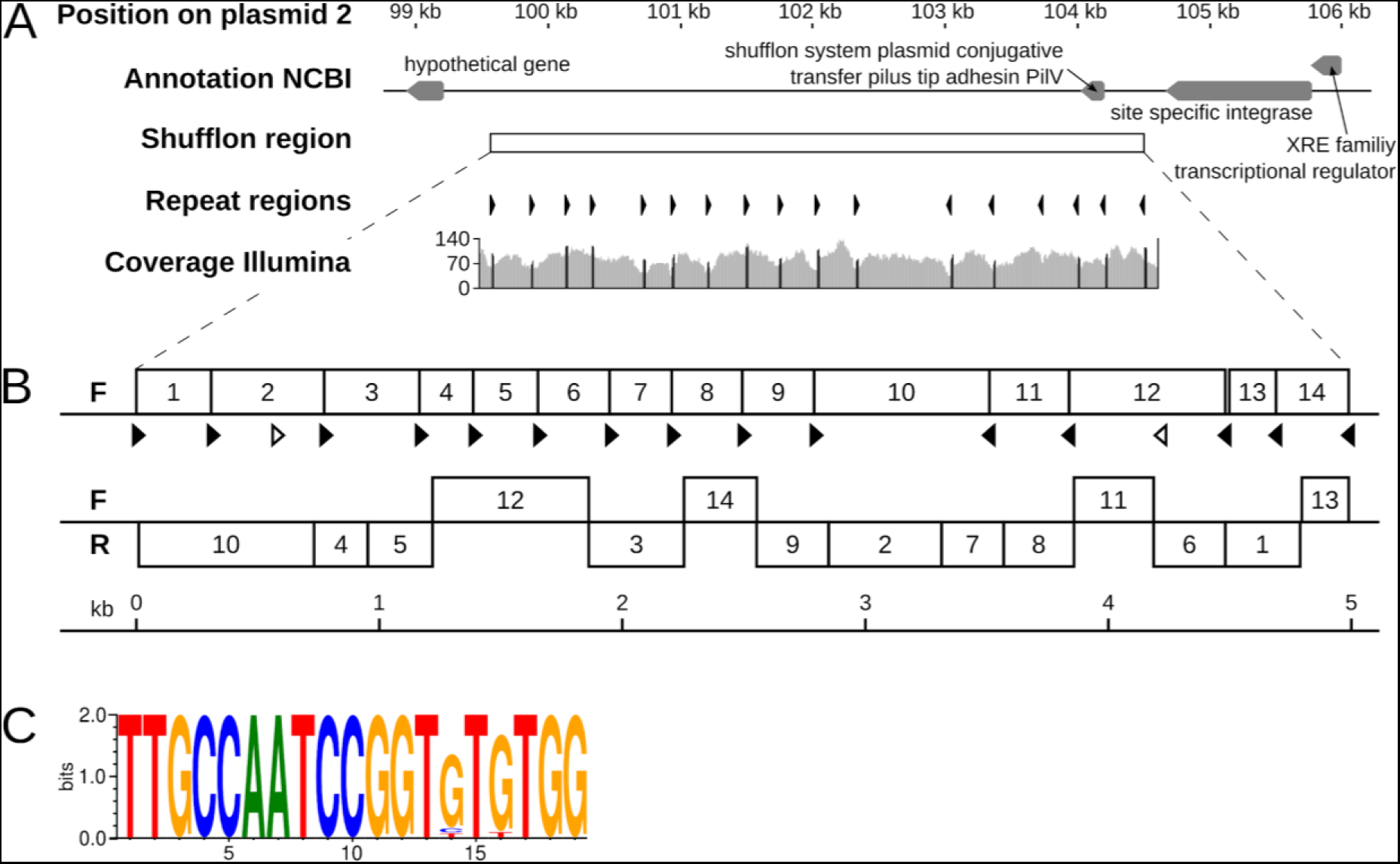
Overview of a shufflon region on plasmid 2. **(A)** The genomic region surrounding the shufflon region on plasmid 2 (top track) is shown along with gene annotations from NCBI (second track). The shufflon region is indicated as white box (third track), and the orientations of the inverted repeats are shown (as black arrows, track 4). The repeats are all pointing inwards to a position at around 102.5 kb. The coverage of mapped Illumina reads is shown (track 5). **(B)** Structure of the two major shufflon compositions identified. The upper version represents the sequence incorporated into the final assembly of P19E3. The lower variant exhibited extensive rearrangement of the shufflon locus (F: forward strand; R: reverse strand). **(C)** Consensus of inverted repeats of 19 bp length represented as a sequence logo (39).

Of note, we detected 19 bp inverted repeats in the shufflon sequence, which all pointed inwards to a position at around 102.5 kb (track 4, Figure 4A). The coverage of mapped Illumina reads (track 5, Figure 4A) exhibited characteristic coverage dips around most repeat positions, with a coverage peak exactly at the repeat position (shown in black on “Coverage Illumina” track, Figure 4A) (49). Importantly, rearrangements for most inverted repeat sites between the two recombination variants were observed (black arrows, Figure 4B), except for two repeat positions which did not show any rearrangement (white arrows). To additionally check the replaced region, ONT and Illumina data were mapped to this region. We could confirm that the mapping in the corrected region was now of a high quality, although showing a lower coverage compared to the rest of plasmid 2, since the mapped reads originated from differently recombined versions of the shufflon. Importantly, a highly conserved sequence motif of the inverted repeats could be identified for this shufflon system (Figure 4C), where only the bases at position 14 and 16 of the 19 bp motif showed variation. The motif sequence was closely related to a repeat reported for an Inc2 plasmid shufflon (49).

### Prevalence of very complex to assemble prokaryotes

To explore the extent of genome assembly complexity for prokaryotes in general, we first analyzed the repeats of 9331 publicly available, complete bacterial genomes. Notably, the existence of complex genomes was not unique for the genus *Pseudomonas*, but a general feature of bacteria. Overall, 663 genomes (7.1%) harbored more than 300 repeats and 300 strains had genomes that contained near identical repeats larger than 30 kb (3.2%) (Figure 5, Supplementary Table S4). A total of 151 genomes were not covered in the plot (all of them bacteria), including 102 genomes with more than 600 repeats and 49 genomes whose longest repeats exceeded 100 kb (Figure 5, upper right box).

**Figure 5.**
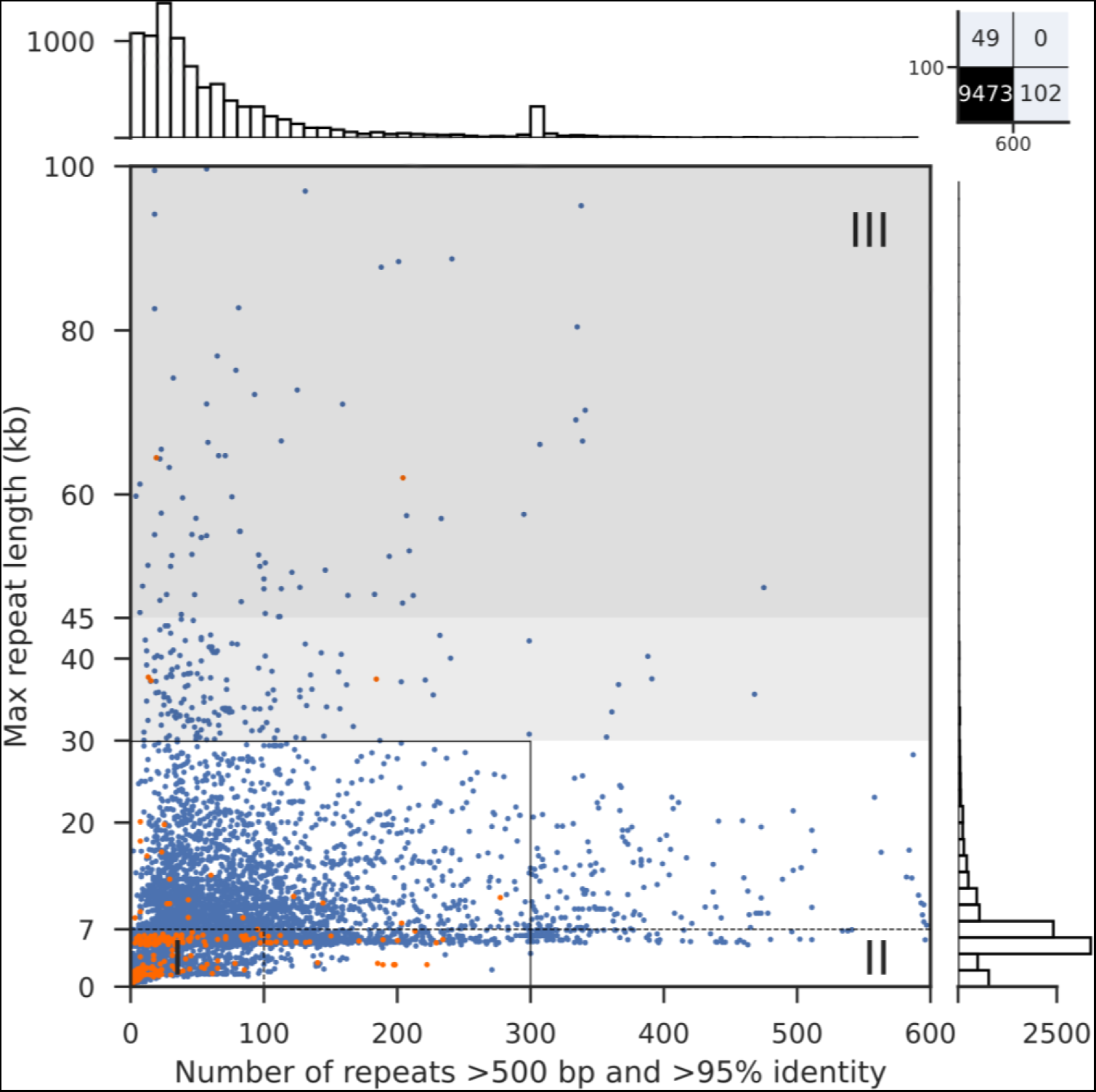
Genome assembly complexity of 9331 bacteria and 293 archaea whose genomes have been completed. Blue/orange dots represent bacterial/archaeal assemblies, respectively, with the overall number of repeats as x-axis coordinate (up to 600 repeats) and the longest repeat as y-axis coordinate (up to 100 kb). The dotted lines separate the three genome assembly complexity classes; the solid black line represents the original plot dimensions by Koren and colleagues (25). The light gray zone represents genomes that potentially can be assembled with PacBio reads depending on the quality of the library and sequencing run. Above 45 kb repeat length (dark gray zone), ONT reads are expected to provide significant benefit whereas closed genomes with PacBio reads only would depend on further advances in technology. Barplots visualize the distribution of the overall number of repeats per strain on top and longest repeat per strain on the right.

Among the complex, overall very repeat-rich genomes the following five species predominated: *Bordetella pertussis* (310 of 663), *Xanthomonas oryzae* (33 of 663), *Yersinia pestis* (31 of 663), *Burkholderia mallei* (28 of 663) and *Enterococcus faecium* (23 of 663). Of note, four of the top species represented pathogens, and the respective percentages of complex class II and class III genomes among the available strains were very high: 89% (310 of 348) for *B. pertussis strains*, 97% (33 of 34) for *X. oryzae* strains, 91% (31 of 34) for *Y. pestis* strains and 100% (28 of 28) for *B. mallei* strains. For the opportunistic pathogen *E. faecium*, 53% (23 of 43) genomes harbored more than 300 repeats. These data might reflect a greater impetus to sequence the full genomes of bacteria with pathogenic lifestyles. In support of this hypothesis, the histogram on top of Figure 5 showed a peak of strains with around 300 repeats. This peak originated from an overrepresentation of *Bordetella pertussis* genomes in NCBI GenBank, apparently due to a focused sequencing project for this pathogen.

Furthermore, for strains whose genomes contained very long near identical repeats, the following five species were most prevalent: *Escherichia coli* (25 of 300), *Francisella tularensis* (14 of 300), *Pseudomonas aeruginosa* (11 of 300), *Burkholderia mallei* (8 of 300) and *Mycobacterium bovis* (7 of 300). For these species, all described as pathogens, only a smaller fraction of the sequenced strains contained many repeats: 10% (11 of 106) for *P. aeruginosa* strains and 5% (25 of 457) for *E. coli.* A higher percentage was observed for *F. tularensis* strains with 38% (14 of 37) and for M. *bovis* strains with 44% (7 of 16). Strikingly, *B. mallei* strains apparently combine both very long repeats (8 of 28, 29%) with a large number of repeats (28 of 28, 100%). They will thus represent a formidable challenge when aiming for a complete *de novo* genome assembly.

We also explored complete genomes of archaea, for which 293 genomes have been deposited at NCBI (Supplementary Table S4). While several genomes harbored more than 200 repeats (7), none of them contained more than 300 repeats. In contrast, 6 genomes featured long repeats above 30 kb (Supplementary Table S4), indicating that a sizeable number of archaea likely also contain very complex genomes.

Finally, for 70 bacterial assemblies we found repeats longer than the 69 kb reported for *P. koreensis* P19E3 (0.7% of 9331 bacterial assemblies; Supplementary Table S4). An examination of 23 of these cases, where the assembly could be easily linked to a publication via its GenBank record, indicated that virtually all of the assemblies had required additional, very labor-intensive methods like cosmid libraries, optical mapping, primer walking or some form of manual curation. In another case, ONT data was used in combination with PacBio to resolve repeats of 212 kb for *Escherichia coli* O157 (see Supplementary Table S4). The longest repeat (2,974,674 bp) was identified for *Calothrix sp.* NIES-4101 (Supplementary Table S4), whose assembly was based on short Illumina reads and with a moderate coverage of 53-fold. A closer inspection revealed that the sequence of plasmid 1 (2,97 Mbp) was entirely contained in the chromosome sequence (7.24 Mbp) and was identical over its entire length (Supplementary Figure S3), suggesting a mis-assembly.

Interestingly, 13 out of the 70 genomes with the longest repeats (18.6%) were from *Streptomyces* species and had linear chromosomes with long terminal inverted repeats (e.g. *Streptomyces pristinaespiralis* HCCB 10218, *Streptomyces sp.* 4F and *Streptomyces lavendulae* subsp. *lavendulae* CCM 3239) (50). Another genome with long terminal inverted repeats (124 kb) was from *Kitasatospora setae* KM-6054, which is closely related to *Streptomyces* (51).

## Discussion

The *de novo* genome assembly of *P. koreensis* P19E3 turned out to be extremely challenging, which could be attributed to the presence of long, near identical repeat pairs of 34, 37, 46 and 69 kb. Only by using very long reads from the ONT platform (mean read length 44 kb) was it possible to fully resolve this genome, indicating the presence of dark matter of genome assembly also in prokaryotes (24). Notably, the long repeats were shared between the chromosome and three large plasmids (Figure 3), a feature we had not observed before in approx. 40 other complete *de novo* assembly projects. Two long, near identical repeats of 69 and 34 kb were located adjacent to each other on the chromosome, only separated by a few bases of unique sequence. This example indicates that the assembly complexity for some genomes may likely even be greater than suggested by mere analysis of individual repeats. In this specific case, an ONT read of 164 kb spanned these two repeats and unique sequences on both ends. Moreover, a highly repetitive shufflon region represented a second major obstacle for complete genome assembly (Figure 4) (52). We could resolve two similarly abundant variants; identification of potential additional, lower abundant variants would require a deeper sequencing of this genomic region, which was beyond the scope of this study.

Importantly, the analysis of all publicly available, complete prokaryotic genomes from NCBI Genbank uncovered an unexpected high percentage of complex genomes. Roughly 7% were class II genomes, which are characterized by a large number or repeats, but none longer than the rDNA operon. For the genus *Pseudomonas*, the plant pathogenic *Psa* strains contained the largest number of repeats (ranging from 277 to 587). We could recently show that such class II genomes can be readily assembled into complete genomes relying on PacBio data (8). In that case, they greatly facilitated a comparative genomics study, as the repeats also contained core genes, that, by relying on Illumina data alone, would have been missed. Similarly, PacBio long reads have facilitated complete *de novo* assembly for many class III genomes (9). However, we here identified a sizeable and particularly difficult subset of strains (roughly 3%) whose genomes harbor long near identical repeats above 30 kb to over 100 kb in length. These cases will greatly benefit from very long ONT reads and appropriate assembly algorithms like Flye (14). As complex genomes will be under-represented *per se* among completely sequenced genomes, we assume that the fraction of complex genomes may even be higher than this estimate. The fraction of class II and III genomes may also increase when results of larger sequencing efforts of diverse prokaryotes like the GEBA initiative (15) and the NCTC 3000 project will become available. So far, we noted a bias for prokaryotes with pathogenic lifestyles, that likely reflects research priorities or funding opportunities. For archaea, more data is needed for a more meaningful comparison with bacteria.

Considering the difficulties in resolving the 69 kb repeats of *P. koreensis* P19E3, even with PacBio and ONT data, the repeat analysis results also raise questions: How were researchers able to assemble genomes with near identical repeats of 100 kb and longer? While only a fraction of the complete genome assemblies specified the sequencing technology they had used, we expect that some of the assemblies that did not rely on PacBio or even ONT data contain assembly errors. In the case of *Pseudomonas aeruginosa* PA1R (53), we noted that the assembly differed in its repeat structure between two assembly versions: while the first assembly relied on Illumina data and contained a 44 kb repeat, the second had relied on PacBio data; there, the repeat was not present any more. Also, the assembly with the longest repeat of 2.97Mb, upon closer inspection, appeared to represent a mis-assembly. We believe that the NCBI would benefit from flagging assemblies with large repeats that are only based on Illumina short read data without evidence for further efforts. A closer inspection of several projects that had assembled genomes with repeats above 70 kb indicated that the researchers had to put in a serious effort: they had either relied on larger insert clone libraries like cosmids (up to about 45 kb inserts), bacterial artificial chromosomes (BACs, with larger insert sizes), primer-walking or some form of manual curation to generate a complete genome sequence. For *P. putida* DLL-E4, the most complex *Pseudomonad*, time-consuming primer-walking was used to close the genome (54), whereas in the case of *P. knackmussi* B13, optical mapping was used (55). With the availability of very long DNA reads and appropriate bioinformatics tools, such labor-intense steps likely will become obsolete for prokaryotes. In our hands, the longest mappable ONT read was 288 kb in length. These advances will further accelerate the exponential increase in complete genome sequences, and will also have a big impact on the *de novo* genome assembly of eukaryotic genomes, as exemplified for the human genome (27), and for nematodes (56).

Finally, we like to caution researchers that plan to rely on a reference-based genome assembly strategy, i.e. mapping sequencing reads against a reference instead of opting for a *de novo* genome assembly: they may easily oversee existing genome differences among closely related strains like single nucleotide variations or larger regions unique to either genome (9). Combined with a publicly available solution for accurate genome annotation of prokaryotes by proteogenomics (9), long reads and modern *de novo* genome assemblers will enable the research community to exploit the increasing number of complete bacterial genome sequences. Amongst other benefits, we expect this will help to unravel new functions encoded by diverse microbiomes, push application of SMRT sequencing for diagnostics (57) and accelerate the identification of urgently needed new antibiotics.

## Acknowledgements

The authors thank Sergej Koren for advice with the repeat analysis, Mikhail Kolmogorov, Jeffrey Yuan, and Pavel A. Pevzner for advice on the assembly of complex genomes with near identical long repeats, F. Freimoser and H.M. Fischer for critical feedback on the manuscript.

## Author contributions

MS and CHA carried out the repeat analysis and devised the overall strategy for genome assembly of complex prokaryotic genomes, MS carried out genome assemblies and identified the structural variation, MRE isolated the *P. koreensis* P19E3 strain and investigated biologically relevant aspects, DF and JFR contributed MiSeq data, AP, RS, DF and JFR contributed ONT long read data, MS and CHA wrote the paper. All authors read and approved the final manuscript.

## Supplementary Tables and Figures

**Supplementary Figure S1.** Overview of workflow to create the final assembly, carry out error correction, and comparison to previous assemblies.

**Supplementary Figure S2.** Maximum likelihood phylogenetic tree placing *P. koreensis* P19E3 in context. P19E3 is shown in bold and it is grouped into a clade of three *P. koreensis* strains. The phylogenetic tree was constructed using amino acid sequence alignment of 107 housekeeping genes. Bootstrap support is shown for all nodes (100 bootstrap runs). The bar at the bottom reflects the number of amino acid changes per site. *Azotobacter vinelandii* DJ served as outgroup. This analysis also contained genomes with assembly level “Chromosome” since this analysis only dealt with phylogenetic aspects, not with assembly complexity. The NCBI GenBank/RefSeq accession numbers are as follows: *Azotobacter vinelandii* DJ: NC_012560; *Pseudomonas brenneri* BS2771: NZ_LT629800; *Pseudomonas citronellolis* P3B5: NZ_CP014158; *Pseudomonas fluorescens* Pf0-1: NC_007492; *Pseudomonas granadensis* LMG 27940: NZ_LT629778; *Pseudomonas koreensis* BS3658: NZ_LT629687; *Pseudomonas koreensis* CRS05-R5: NZ_CP015852; *Pseudomonas koreensis* D26: NZ_CP014947; *Pseudomonas koreensis* P19E3: CP027477; *Pseudomonas mandelii* LMG 21607: NZ_LT629796; *Pseudomonas moraviensis* BS3668: NZ_LT629788; *Pseudomonas putida* W619: NC_010501; *Pseudomonas reinekei* BS3776: NZ_LT629709; *Pseudomonas* sp. B10: NZ_LT707063; *Pseudomonas* sp. DR 5-09: NZ_CP011566; *Pseudomonas* sp. Z003-0.4C(8344-21): NZ_LT629756; *Pseudomonas stutzeri* A1501: NC_009434; *Pseudomonas syringae* pv. *tomato* DC3000: NC_004578.

**Supplementary Table S3.** A Gepard dotplot (Krumsiek,J., Arnold,R. and Rattei,T. (2007) Gepard: a rapid and sensitive tool for creating dotplots on genome scale. *Bioinformatics*, **23**, 1026–1028.) showing the alignment of the chromosome (AP018280.1) and plasmid 1 (AP018274.1) of the genome assembly of *Calothrix sp.* NIES-4101. Of note, the sequence of plasmid 1 has 100.0% sequence identity to the end of the chromosome, but is the reverse complement.

**Supplementary Table S1.** Overview of the most repeat-rich *Pseudomonas* genomes. Twelve strains with the highest overall number of repeats are shown.

**Supplementary Table S2.** Different assembly stages compared to the final Illumina-polished assembly.

**Supplementary Table S3.** Mapping statistics of reads from all three sequencing technologies to the final Illumina-polished assembly.

**Supplementary Table S4**. Repeat analysis overview for 9331 bacterial and 293 archaeal complete genomes. The classification for genome assembly complexity is also given. Genomes without repeats longer than 500 bp and a similarity higher than 95% were binned and listed as having no repeats. Genomes are sorted in descending order based on their longest repeat. We kindly ask users to reference our paper.

## Data access

| Database | Accession number(s) |
| --- | --- |
| NCBI GenBank | CP027477 to CP027481 |
| NCBI BioProject | PRJNA436895 |
| NCBI BioSample | SAMN08634678 |

